# Genome-wide association study results for educational attainment aid in identifying genetic heterogeneity of schizophrenia

**DOI:** 10.1101/114405

**Authors:** V. Bansal, M. Mitjans, C.A.P. Burik, R.K. Linnér, A. Okbay, C.A. Rietveld, M. Begemann, S. Bonn, S. Ripke, R. de Vlaming, M.G. Nivard, H. Ehrenreich, P.D. Koellinger

**Author notes:** These co-authors contributed equally (V. Bansal, M. Mitjans). These authors jointly directed this work (M.G. Nivard, H. Ehrenreich, P.D. Koellinger). Correspondence: Professor P.D. Koellinger, Vrije Universiteit Amsterdam, Department of Economics, De Boelelaan 1105, 1081 HV, Amsterdam, Netherlands.

## Abstract

Higher educational attainment (EA) is negatively associated with schizophrenia (SZ). However, recent studies found a positive genetic correlation between EA and SZ. We investigated possible causes of this counterintuitive finding using genome-wide association study results for EA and SZ (N = 443,581) and a replication cohort (1,169 controls; 1,067 cases) with deeply phenotyped SZ patients. We found strong genetic dependence between EA and SZ that cannot be explained by chance, linkage disequilibrium, or assortative mating. Instead, several genes seem to have pleiotropic effects on EA and SZ, but without a clear pattern of sign concordance. Genetic heterogeneity of SZ contributes to this finding. We demonstrate this by showing that the polygenic prediction of clinical SZ symptoms can be improved by taking the sign concordance of loci for EA and SZ into account. Furthermore, using EA as a proxy phenotype, we isolate *FOXO6* and *SLITRK1* as novel candidate genes for SZ.

## MAIN TEXT

Schizophrenia (SZ) is the collective term used for a severe, highly heterogeneous and costly psychiatric disorder that is caused by environmental and genetic factors^1–4^. A genome-wide association study (GWAS) by the Psychiatric Genomics Consortium (PGC) identified 108 genomic loci that are associated with SZ^5^. These 108 loci jointly account for ≈3.4% of the variation on the liability scale for SZ^5^, while all single nucleotide polymorphisms (SNPs) that are currently measured by SNP arrays capture ≈64% (s.e. = 8%) of the variation in liability for the disease^6^. This implies that many genetic variants with small effect sizes contribute to the heritability of SZ, but most of them are unidentified as of yet. A polygenic score (PGS) based on all SNPs currently accounts for 4-15% of the variation on the liability scale for SZ^5^.

However, this PGS does not predict any differences in symptoms or severity of the disease among SZ patients^4^. Partly, this could be because the clinical disease classification of SZ spans several different behavioural and cognitive traits that may not have identical genetic architectures. Therefore, identifying additional genetic variants and understanding through which pathways they are linked with the clinical diagnosis of SZ is an important step in understanding the aetiologies of the ‘schizophrenias’^7^. However, GWAS analyses of specific SZ symptoms would require very large sample sizes to be statistically well powered, and the currently available datasets on deeply phenotyped SZ patients are not yet large enough for this purpose.

Here, we use an alternative approach to make progress with data that is readily available – by combining GWAS for SZ and educational attainment (EA). The GWAS sample sizes for EA are the largest to date for any cognition-related phenotype. Furthermore, previous studies suggest a complex relationship between EA and SZ^8, 9^ that may be used to gain additional insights into the genetic architecture of SZ and its symptoms. In particular, phenotypic data seem to suggest a *negative* correlation between EA and SZ^10^. For example, SZ patients with lower EA typically show an earlier age of disease onset, higher levels of psychotic symptomatology, and worsened global cognitive function^10^. In fact, EA has been suggested to be a measure of premorbid function and a predictor of outcomes in SZ. Moreover, it has been forcefully argued that retarded intellectual development, global cognitive impairment during childhood, and bad school performance should be seen as core features of SZ that precede the development of psychotic symptoms and differentiate SZ from bipolar disorder (BIP)^11–15^. Furthermore, credible genetic links between SZ and impaired cognitive performance have been found^16^.

In contrast to these findings, recent studies using large-scale GWAS results identified a small, but *positive genetic* correlation between EA and SZ (*ρ_EA,SZ_* = 0.08)^8^, and higher PGS values for SZ have been reported to be associated with creativity and greater EA^17^. Other statistically well-powered studies found that a high intelligence quotient (IQ) has protective effects against SZ^18^ and reported a negative genetic correlation between IQ and SZ (*ρ_IQ,SZ_* = −0.2)^19^, suggesting the possibility that genetic effects that contribute to EA *but not via IQ* are responsible for the observed positive genetic correlation between SZ and EA.

Indeed, previous research by the Social Science Genetic Association Consortium (SSGAC)^8^ already demonstrated that the effect of the EA-PGS on years of schooling is mediated by several individual characteristics that have imperfect or no genetic correlation with each other, including higher IQ, higher openness, and higher conscientiousness. These different factors that contribute to EA seem to be related to SZ and its symptoms in complex ways^20–22^. For example, differences in openness have been reported to differentiate between patients diagnosed with schizophrenia spectrum personality disorders (higher openness) from patients diagnosed with SZ (lower openness), while conscientiousness tends to be reduced among patients of both disorders compared to healthy controls^20^.

The contributing factors to EA that have previously been identified by the SSGAC (i.e. IQ, openness, and conscientiousness)^8^ are phenotypically and genetically related, but by no means identical^23, 24^. Specifically, the Cognitive Genomics Consortium (COGENT) reported a moderate genetic correlation between IQ and openness (*r_g_* = 0.48, *P* = 3.25 × 10^−4^), but only a small genetic correlation of IQ and conscientiousness of 0.10 that was indistinguishable from zero (*r_g_* = 0.10, *P* = 0.46)^25^. Therefore, it is appropriate to think of EA as a genetically heterogeneous trait that can be decomposed into subphenotypes that have imperfect genetic correlations with each other. If the various symptoms of SZ also have nonidentical genetic architectures, this could result in a pattern where both EA and SZ share many genetic loci, but without a clear pattern of sign concordance and with seemingly contradictory phenotypic and genetic correlation results.

To explore this hypothesis and to discern it from alternative explanations, we performed a series of statistical genetic analyses using large-scale GWAS results for SZ and EA from nonoverlapping samples. We started by characterizing the genetic relationship between both traits by using EA as a “proxy phenotype”^26^ for SZ. We annotated possible biological pathways, tissues, and cell types implied by genetic variants that are associated with both traits and explored to what extent these variants are also enriched for association with other traits. We tested if the genetic relationship between EA and SZ can be explained by chance, linkage disequilibrium (LD), or assortative mating. Furthermore, we investigated the hypothesis that the part of SZ that is different from BIP is a neurodevelopmental disorder, whereas the part of SZ that overlaps with BIP is not. Finally, we developed a formal statistical test for genetic heterogeneity of SZ using a polygenic prediction framework that leverages both the SZ and the EA GWAS results.

## RESULTS

As a formal prelude to our study, it is conceptually important to differentiate between genetic dependence and genetic correlation. In our analyses, genetic dependence means that the genetic variants associated with EA are more likely to also be associated with SZ than expected by chance. In contrast, genetic correlation is defined by the correlation of the (true) effect sizes of genetic variants on the two traits. Thus, genetic correlation implies a linear genetic relationship between two traits whereas genetic dependence does not. Thus, two traits can be genetically dependent even if they are not genetically correlated and *vice versa.* One possible cause of a non-linear genetic dependence is that at least one of the traits is genetically heterogeneous in the sense that it aggregates across subphenotypes (or symptoms) with non-identical genetic architectures. **Supplementary Note 1** presents a more formal discussion and simulations that illustrate the data patterns that can emerge.

### Proxy-phenotype analyses

We used the proxy-phenotype method (PPM)^26^ to illustrate the genetic dependence between EA and SZ. PPM is a two-stage approach. In the first stage, a GWAS on the proxy-phenotype (EA) is conducted. The most strongly associated loci are then advanced to the second stage, which tests the association of these loci with the phenotype of interest (SZ) in an independent sample. If the two traits are genetically dependent, this two-stage approach can increase the statistical power for detecting associations for the target trait because it limits the multiple testing burden for the phenotype of interest compared to a GWAS^8, 9, 26^.

Our PPM analyses followed a preregistered analysis plan (https://osf.io/dnhfk/) using GWAS results on EA (n = 363,502)^8^ and SZ (34,409 cases and 45,670 controls)^27^ that were obtained from non-overlapping samples of Europeans. For replication and follow-up analyses, we used the Göttingen Research Association for Schizophrenia (GRAS) data collection^28^, which has a uniquely rich and accurate set of SZ measures. The GRAS sample was not part of either GWAS (**Supplementary Notes 2-3**).

Analyses were performed using 8,240,280 autosomal SNPs that passed quality controls in both GWAS and additional filters described in Methods and **Supplementary Note 4**. We selected approximately independent lead SNPs from the EA GWAS that passed the predefined significance threshold of P_EA_ < 10^−5^ and looked up their SZ results. To test if EA-associated SNPs are more strongly associated with SZ than expected by chance (referred to as “raw enrichment” below), we conducted a Mann-Whitney test that compares the *Psz-* values of the EA-associated lead SNPs with the *Psz*-values of a set of randomly drawn, approximately LD-independent SNPs with similar minor allele frequencies (**Supplementary Notes 5-6**). **Fig. 1** presents an overview of the proxy-phenotype analyses.

**Figure 1:**
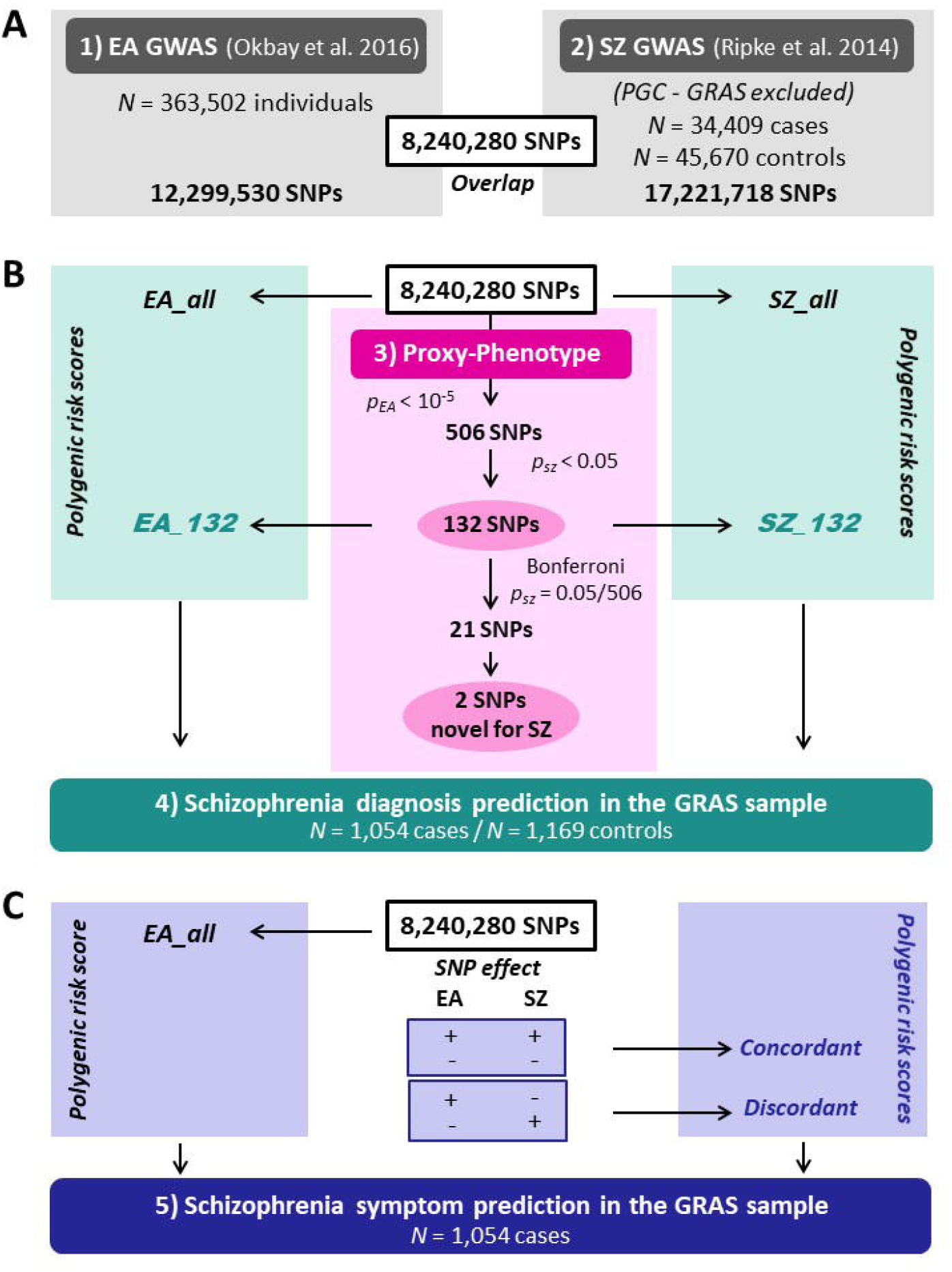
Workflow of the proxy-phenotype analyses. *Notes*: Educational attainment (EA) and schizophrenia (SZ) GWAS results are based on the analyses reported in ref.^5, 8^. All cohorts that were part of the SZ GWAS were excluded from the meta-analysis on EA. The GRAS data collection was not included in either the SZ or the EA meta-analysis. Proxy-phenotype analyses were conducted using 8,240,280 autosomal SNPs that passed quality control. Genetic outliers of non-European descent (*N* = 13 cases) were excluded from the analysis in the GRAS data collection.

The first-stage GWAS on EA (**Supplementary Note 2**) identified 506 loci that passed our predefined threshold of P_EA_ <10^−5^; 108 of them were significant at the genome-wide level (P_EA_ < 5 × 10^−8^, see **Supplementary Table 2**). Of the 506 EA lead-SNPs, 132 are associated with SZ at nominal significance (P_SZ_ < 0.05), and 21 of these survive Bonferroni correction 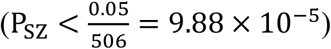 (**Table 1**). LD score regression results suggest that the vast majority of the association signal in both the EA^8^ and the SZ^5^ GWAS are truly genetic signals, rather than spurious signals originating from uncontrolled population stratification. Figure 2a shows a Manhattan plot for the GWAS on EA highlighting SNPs that were also significantly associated with SZ (turquoise crosses for P_SZ_ < 0.05, magenta crosses for P_SZ_ = 9.88 × 10^−5^).

**Figure 2:**
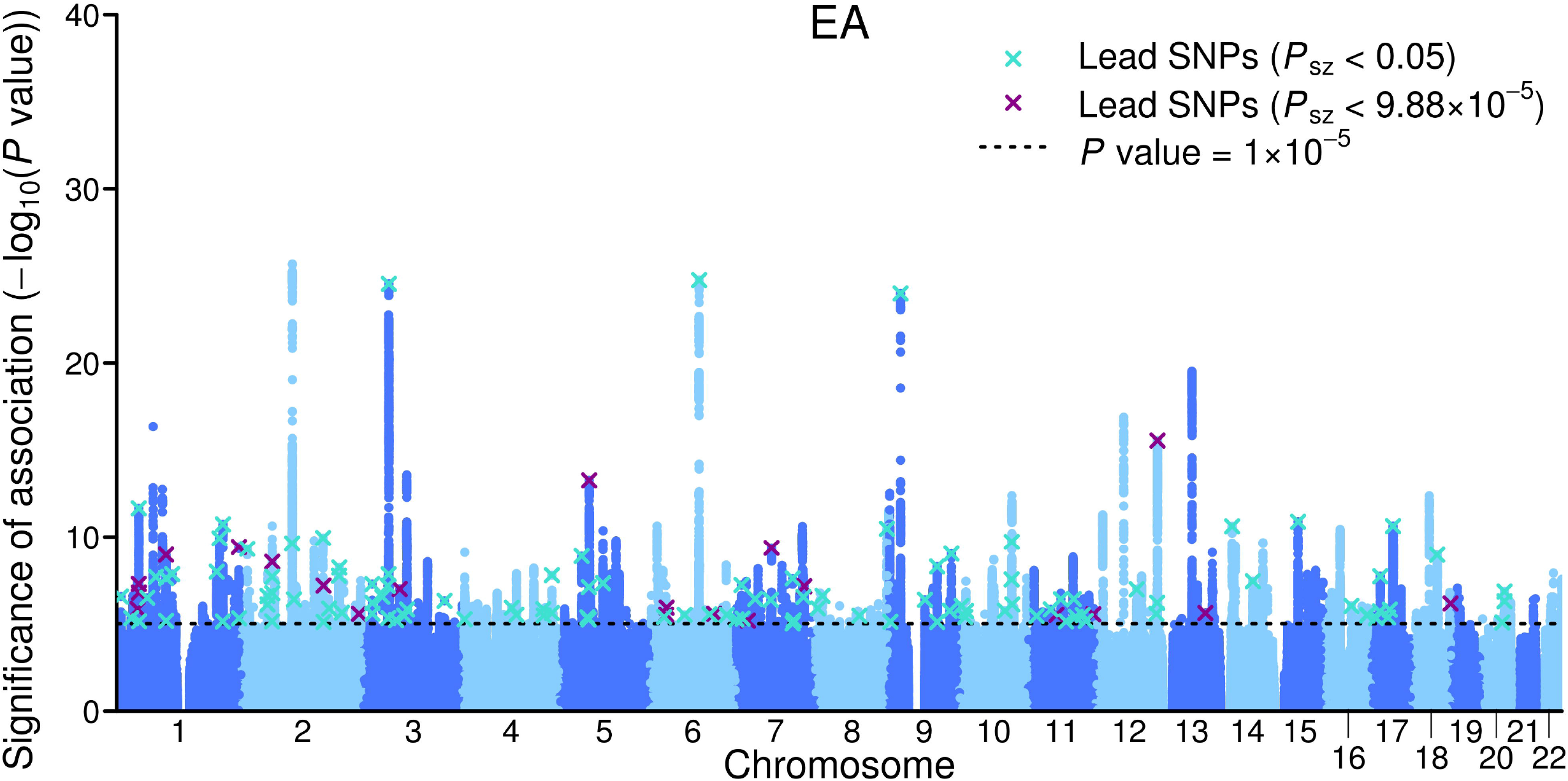
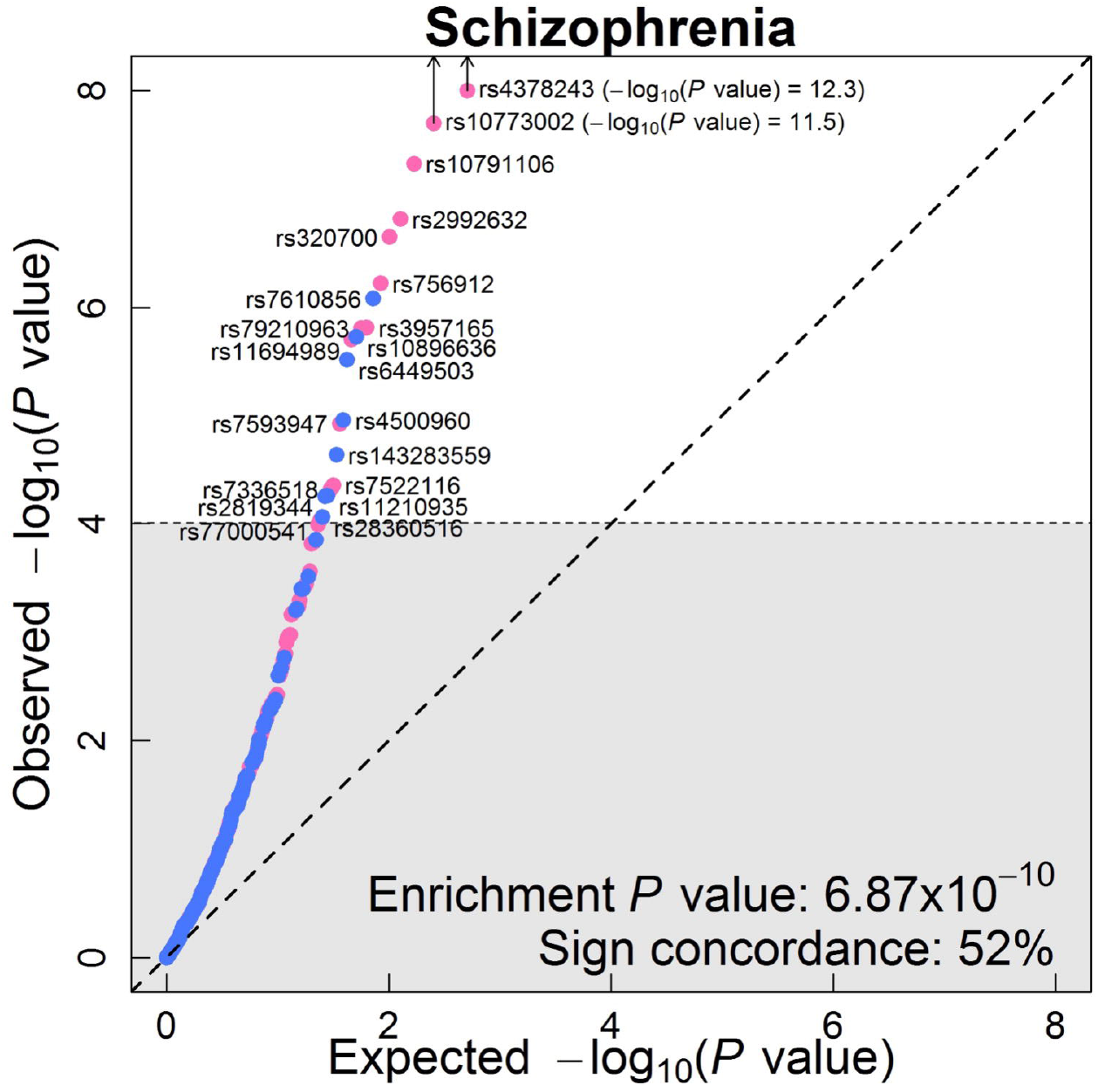
Results of the proxy-phenotype analyses. *Notes:* Panel a: **Manhattan plot for educational attainment (EA) associations (*n* = 363,502)**. The x-axis is the chromosomal position, and the y-axis is the significance on a −log10 scale (2-sided). The black dashed line shows the suggestive significance level of 10^−5^ that we specified in our preregistered analysis plan. Turquois and magenta crosses identify EA-associated lead-SNPs that are also associated with SZ at nominal or Bonferroni-adjusted significance levels, respectively. Panel b: **Q–Q plot of the 506 EA-associated SNPs for schizophrenia (SZ) (n = 34,409 cases and *n* = 45,670 controls)**. SNPs with concordant effects on both phenotypes are pink, and SNPs with discordant effects are blue. SNPs outside the grey area (21 SNPs) pass the Bonferroni-corrected significance threshold that corrects for the total number of SNPs we tested (*P* < 0.05/506 = 9.88×10^−5^) and are labelled with their rs numbers. Observed and expected *P* values are on a −log10 scale. For the sign concordance test: *P* = 0.40, 2-sided.

**Table 1:**
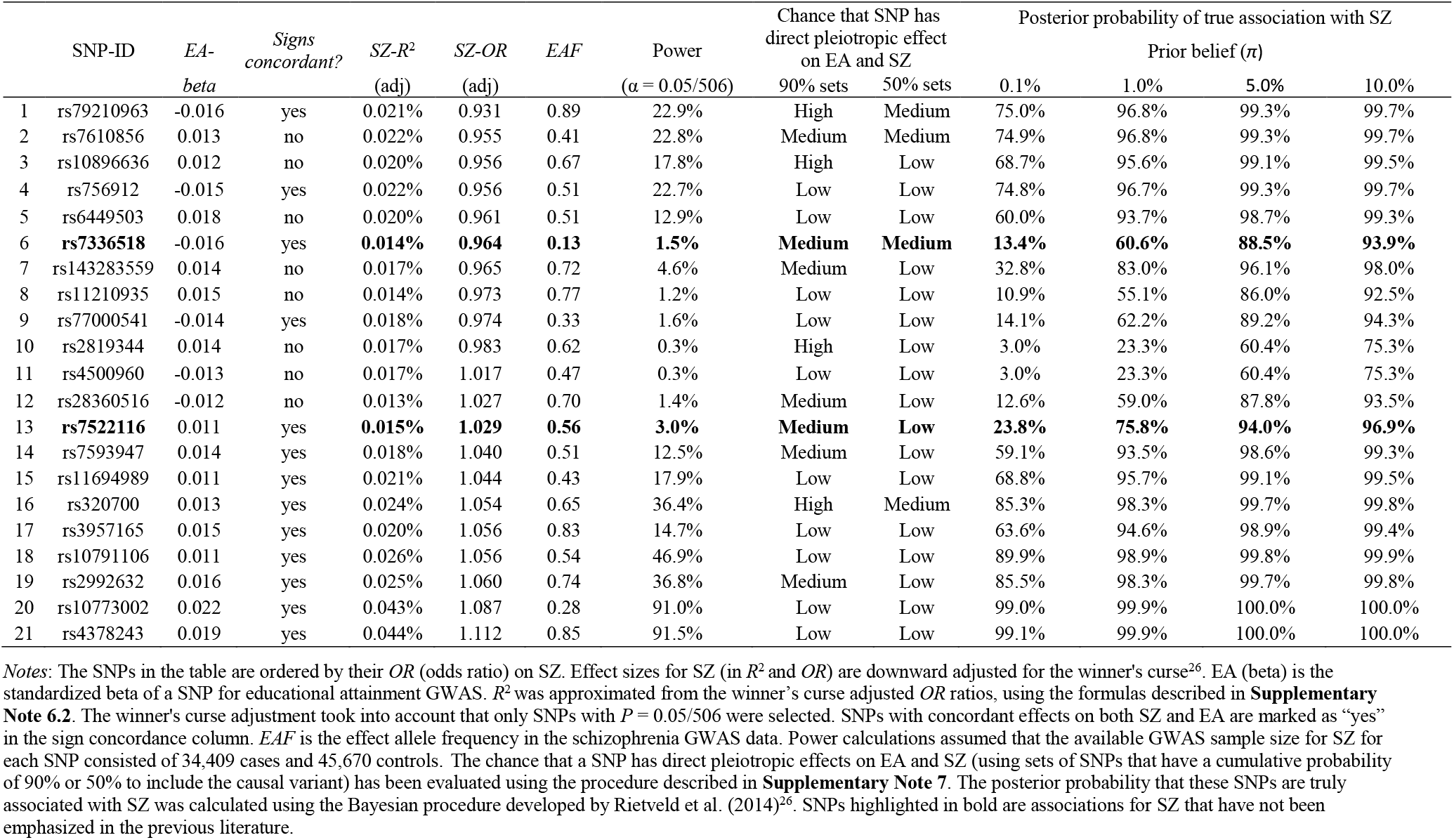
SNPs significantly associated with schizophrenia after Bonferroni correction.

| | SNP-ID | EA-<br>*beta* | *Signs*<br>*concordant?* | SZ-$R^2$<br>(adj) | SZ-OR<br>(adj) | EAF | Power<br>($\alpha = 0.05/506$) | Chance that SNP has<br>direct pleiotropic effect<br>on EA and SZ | | Posterior probability of true association with SZ | | | |
| --- | --- | --- | --- | --- | --- | --- | --- | --- | --- | --- | --- | --- | --- |
| | | | | | | | | 90% sets | 50% sets | Prior belief ($\pi$) | | | |
|  |  |  |  |  |  |  |  |  |  | 0.1% | 1.0% | 5.0% | 10.0% |
| 1 | rs79210963 | -0.016 | yes | 0.021% | 0.931 | 0.89 | 22.9% | High | Medium | 75.0% | 96.8% | 99.3% | 99.7% |
| 2 | rs7610856 | 0.013 | no | 0.022% | 0.955 | 0.41 | 22.8% | Medium | Medium | 74.9% | 96.8% | 99.3% | 99.7% |
| 3 | rs10896636 | 0.012 | no | 0.020% | 0.956 | 0.67 | 17.8% | High | Low | 68.7% | 95.6% | 99.1% | 99.5% |
| 4 | rs756912 | -0.015 | yes | 0.022% | 0.956 | 0.51 | 22.7% | Low | Low | 74.8% | 96.7% | 99.3% | 99.7% |
| 5 | rs6449503 | 0.018 | no | 0.020% | 0.961 | 0.51 | 12.9% | Low | Low | 60.0% | 93.7% | 98.7% | 99.3% |
| 6 | **rs7336518** | -0.016 | yes | **0.014%** | **0.964** | **0.13** | **1.5%** | **Medium** | **Medium** | **13.4%** | **60.6%** | **88.5%** | **93.9%** |
| 7 | rs143283559 | 0.014 | no | 0.017% | 0.965 | 0.72 | 4.6% | Medium | Low | 32.8% | 83.0% | 96.1% | 98.0% |
| 8 | rs11210935 | 0.015 | no | 0.014% | 0.973 | 0.77 | 1.2% | Low | Low | 10.9% | 55.1% | 86.0% | 92.5% |
| 9 | rs77000541 | -0.014 | yes | 0.018% | 0.974 | 0.33 | 1.6% | Low | Low | 14.1% | 62.2% | 89.2% | 94.3% |
| 10 | rs2819344 | 0.014 | no | 0.017% | 0.983 | 0.62 | 0.3% | High | Low | 3.0% | 23.3% | 60.4% | 75.3% |
| 11 | rs4500960 | -0.013 | no | 0.017% | 1.017 | 0.47 | 0.3% | Low | Low | 3.0% | 23.3% | 60.4% | 75.3% |
| 12 | rs28360516 | -0.012 | no | 0.013% | 1.027 | 0.70 | 1.4% | Medium | Low | 12.6% | 59.0% | 87.8% | 93.5% |
| 13 | **rs7522116** | 0.011 | yes | **0.015%** | **1.029** | **0.56** | **3.0%** | **Medium** | **Low** | **23.8%** | **75.8%** | **94.0%** | **96.9%** |
| 14 | rs7593947 | 0.014 | yes | 0.018% | 1.040 | 0.51 | 12.5% | Medium | Low | 59.1% | 93.5% | 98.6% | 99.3% |
| 15 | rs11694989 | 0.011 | yes | 0.021% | 1.044 | 0.43 | 17.9% | Low | Low | 68.8% | 95.7% | 99.1% | 99.5% |
| 16 | rs320700 | 0.013 | yes | 0.024% | 1.054 | 0.65 | 36.4% | High | Medium | 85.3% | 98.3% | 99.7% | 99.8% |
| 17 | rs3957165 | 0.015 | yes | 0.020% | 1.056 | 0.83 | 14.7% | Low | Low | 63.6% | 94.6% | 98.9% | 99.4% |
| 18 | rs10791106 | 0.011 | yes | 0.026% | 1.056 | 0.54 | 46.9% | Low | Low | 89.9% | 98.9% | 99.8% | 99.9% |
| 19 | rs2992632 | 0.016 | yes | 0.025% | 1.060 | 0.74 | 36.8% | Medium | Low | 85.5% | 98.3% | 99.7% | 99.8% |
| 20 | rs10773002 | 0.022 | yes | 0.043% | 1.087 | 0.28 | 91.0% | Low | Low | 99.0% | 99.9% | 100.0% | 100.0% |
| 21 | rs4378243 | 0.019 | yes | 0.044% | 1.112 | 0.85 | 91.5% | Low | Low | 99.1% | 99.9% | 100.0% | 100.0% |

*Notes:* The SNPs in the table are ordered by their $OR$ (odds ratio) on SZ. Effect sizes for SZ (in $R^2$ and $OR$ ) are downward adjusted for the winner's curse<sup>26</sup>. EA ( $\beta$ ) is the standardized $\beta$ of a SNP for educational attainment GWAS. $R^2$ was approximated from the winner's curse adjusted $OR$ ratios, using the formulas described in **Supplementary Note 6.2**. The winner's curse adjustment took into account that only SNPs with $P = 0.05/506$ were selected. SNPs with concordant effects on both SZ and EA are marked as "yes" in the sign concordance column. $EAF$ is the effect allele frequency in the schizophrenia GWAS data. Power calculations assumed that the available GWAS sample size for SZ for each SNP consisted of 34,409 cases and 45,670 controls. The chance that a SNP has direct pleiotropic effects on EA and SZ (using sets of SNPs that have a cumulative probability of 90% or 50% to include the causal variant) has been evaluated using the procedure described in **Supplementary Note 7**. The posterior probability that these SNPs are truly associated with SZ was calculated using the Bayesian procedure developed by Rietveld et al. (2014)<sup>26</sup>. SNPs highlighted in bold are associations for SZ that have not been emphasized in the previous literature.

A Q-Q plot of the 506 EA lead SNPs for SZ is shown in **Figure 2b**. Although the observed sign concordance of 52% is not significantly different from a random pattern (*P* = 0.40), we find 3.23 times more SNPs in this set of 506 SNPs that are nominally significant for SZ than expected given the distribution of the *P* values in the SZ GWAS results (raw enrichment P = 6.87 × 10^−10^, **Supplementary Note 6**). The observed enrichment of the 21 EA lead SNPs that pass Bonferroni correction for SZ 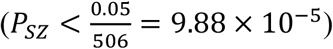 is even more pronounced (27 times stronger, *P* = 5.44 × 10^−14^).

**Table 2:** Polygenic prediction of schizophrenia measures in the GRAS patient sample.

|  |  | Years of<br>education<sup>1</sup> | Age at<br>prodrome | Age at disease<br>onset | Premorbid IQ<sup>1</sup> | GAF<sup>2</sup> | CGI-S<sup>2</sup> | PANSS<br>positive<sup>2</sup> | PANSS<br>negative<sup>2</sup> |
| --- | --- | --- | --- | --- | --- | --- | --- | --- | --- |
| **Baseline Model** |  |  |  |  |  |  |  |  |  |
| *SZ_all* | *standardized beta* | 0.001 | -0.041 | -0.056 | -0.063 | -0.024 | 0.041 | 0.033 | 0.043 |
|  | *P value* | 0.976 | 0.297 | 0.129 | 0.090 | 0.510 | 0.249 | 0.364 | 0.253 |
| *EA_all* | *standardized beta* | 0.182** | 0.005 | -0.002 | 0.149** | 0.068* | -0.057 | 0.001 | -0.051 |
|  | *P value* | 4.4x10<sup>-09</sup> | 0.884 | 0.961 | 7.2x10<sup>-06</sup> | 0.029 | 0.065 | 0.981 | 0.107 |
|  | *Adj. R<sup>2</sup>* | 0.0612 | 0.0023 | 0.0047 | 0.0417 | 0.0655 | 0.0816 | 0.0711 | 0.0243 |
| | $\Delta$ *Adj. R<sup>2</sup>*# | 0.0312 | -0.0010 | -0.0009 | 0.0209 | 0.0035 | 0.0023 | -0.0010 | 0.0015 |
| **Split Model** |  |  |  |  |  |  |  |  |  |
| **Concordant** | *standardized beta* | -0.013 | -0.019 | -0.031 | -0.043 | -0.096* | 0.050 | 0.079 | 0.125** |
|  | *P value* | 0.751 | 0.665 | 0.456 | 0.326 | 0.022 | 0.232 | 0.059 | 0.0036 |
| **Discordant** | *standardized beta* | 0.014 | -0.030 | -0.035 | -0.034 | 0.066 | <0.001 | -0.039 | -0.072 |
|  | *P value* | 0.730 | 0.515 | 0.409 | 0.437 | 0.112 | 0.996 | 0.351 | 0.090 |
| *EA_all* | *standardized beta* | 0.191** | 0.002 | -0.002 | 0.153** | 0.122** | -0.074 | -0.039 | -0.118** |
|  | *P value* | 1.0x10<sup>-06</sup> | 0.965 | 0.953 | 2.7x10<sup>-04</sup> | 0.002 | 0.058 | 0.319 | 0.003 |
|  | *Adj. R<sup>2</sup>* | 0.0604 | 0.0012 | 0.0037 | 0.0406 | 0.0694 | 0.0811 | 0.0728 | 0.0306 |
| | $\Delta$ *Adj. R<sup>2</sup>*# | 0.0304 | -0.0021 | -0.0019 | 0.0198 | 0.0074 | 0.0018 | 0.0007 | 0.0078 |
|  | *n* | 1,039 | 915 | 1,043 | 903 | 1,010 | 1,014 | 1,009 | 1,002 |
| | $\Delta R^2$ (*Split Model* – *Baseline Model*) | -0.0008 | -0.0011 | -0.0010 | -0.0011 | 0.0039 | -0.0005 | 0.0017 | 0.0063 |
|  | *P value from F-test*<sup>o</sup> | 0.698 | 0.907 | 0.968 | 0.891 | 0.023* | 0.479 | 0.098 | 0.007** |

Notes: Linear regression using the first 10 genetic principal components as control variables. <sup>1</sup>: Age of onset was included as covariate. <sup>2</sup>: Medication was included as covariate. # Change in *Adj. R<sup>2</sup>* of the models compared to a model that only contains the *SZ\_all* score and the control variables. <sup>o</sup> *P value* from *F-test* refers to improvement in split model compared to baseline model. \*denotes significance at $P < 0.05$ . \*\*denotes significance at $P < 0.01$ .

The effect sizes of these 21 SNPs on SZ are small, ranging from Odds Ratio (*OR*) = 1.02 (rs4500960) to *OR* = 1.11 (rs4378243) after correction for the statistical winner’s curse^26^ (**Table 1**). We calculated the probability that these 21 SNPs are truly associated with SZ using a heuristic Bayesian method that takes the winner’s curse corrected effect sizes, statistical power, and prior beliefs into account^26^. Applying a reasonable prior belief of 5% (**Supplementary Note 6**), we find that all 21 SNPs are likely or almost certain to be true positives.

### Prediction of future genome-wide significant loci for schizophrenia

Of the 21 variants we identified, 12 are in LD with loci previously reported by the PGC^5^ and 2 are in the major histocompatibility complex region on chromosome (chr) 6 and were therefore not separately reported in that study. Three of the variants we isolated (rs7610856, rs143283559, and rs28360516) were independently found in a meta-analysis of the PGC results^5^ with another large-scale sample^29^. We show in **Supplementary Note 6** that using EA as a proxy-phenotype for SZ helped to predict the novel genome-wide significant findings reported in that study, which illustrates the power of the proxy-phenotype approach. Furthermore, two of the 21 variants (rs756912, rs7593947) are in LD with loci recently reported in a study that also compared GWAS findings from EA and SZ using smaller samples and a less conservative statistical approach^30^. The remaining 2 SNPs we identified (rs7336518 on chr13 and rs7522116 on chr1) add to the list of empirically plausible candidate loci for SZ.

### Detection of shared causal loci

The next step in our study was a series of analyses that aimed to identify reasons for the observed genetic dependence between EA and SZ and to put the findings of the PPM analysis into context. First, we probed if there is evidence that the loci identified by the PPM may tag shared causal loci for both EA and SZ (i.e., pleiotropy), rather than being in LD with different causal loci for both traits.

For each of the 21 SNPs isolated by our PPM analysis, we looked at their neighboring SNPs within a +/-500 kb window and estimated their posterior probability of being causal for EA or SZ using PAINTOR^31^. We then selected two sets of SNPs, each of which contains the smallest number of SNPs that yields a cumulative posterior probability of 90% or 50% of containing the causal locus for EA and SZ. We refer to these as broad sets (90%) and narrow sets (50%), respectively. (**Supplementary Note 7** also describes results for the 80% and 65% credibility sets). For each of these sets, we calculated the posterior probability that it contains the causal locus for the other trait. We classify the probability of a locus being pleiotropic as low, medium, or high if the posterior probability of both the EA set on SZ and the SZ set on EA are <15%, 15%-45%, or >45%, respectively (**Supplementary Note 7**).

For the broad credibility set analyses (90%), we found eleven loci with a medium or high credibility to have direct causal effects on both EA and SZ (including one of the novel SNPs, rs7336518). Six of these loci have concordant effects on the two traits (i.e., ++ or --) while five have a discordant effects (i.e., +- or -+, **Table 1** and **Supplementary Note 7**). Overall, our analyses suggest that some of the 21 SNPs that we identified by using EA as a proxy-phenotype for SZ are likely to have direct pleiotropic effects on both traits. Of the most likely candidates for direct pleiotropic effects, three SNPs have concordant signs (rs79210963, rs7336518, rs320700) and one has discordant signs (rs7610856).

### Biological annotations

Biological annotation of the 132 SNPs that are jointly associated with EA (*P_EA_* < 10^−5^) and SZ (*P_sz_* < 0.05) using DEPICT (**Supplementary Note 8**) points to genes that are known to be involved in neurogenesis and synapse formation (**Supplementary Table 4.4**). Some of the indicated genes, including *SEMA6D* and *CSPG5,* have been suggested to play a potential role in SZ^32, 33^. For the two novel candidate SNPs reported in this study (rs7522116 and rs7336518), DEPICT points to the *FOXO6* (Forkhead Box O6) and the *SLITRK1* (SLIT and NTRK Like Family Member 1) genes, respectively. *FOXO6* is predominantly expressed in the hippocampus and has been suggested to be involved in memory consolidation, emotion and synaptic function^34, 35^. Similarly, *SLITRK1* is also highly expressed in the brain^36^, is particularly localized to excitatory synapses and promotes their development^37^, and it has previously been suggested to be a candidate gene for neuropsychiatric disorders^38^.

### LD-aware enrichment across different traits

The raw enrichment *P* value reported in Fig. 2b and Supplementary Note 6 could in principle be due to the LD-structure of the EA lead SNPs that we tested. Specifically, if these EA lead SNPs have stronger LD with other SNPs in the human genome than expected by chance, this could cause the observed enrichment of this set of SNPs on SZ and other traits because higher LD increases the chance these SNPs would “tag” causal SNPs that they are correlated with^39, 40^.

To assess the null hypothesis that the observed genetic dependence between EA and SZ can be entirely explained by LD patterns in the human genome, we developed an association enrichment test that corrects for the LD score of each SNP (**Supplementary Note 9**). We applied this test to the 132 SNPs that are jointly associated with EA (*P_EA_* < 10^−5^) and SZ (*P_SZ_* < 0.05), i.e. the loci that were identified by using EA as a proxy-phenotype for SZ. LD scores were obtained from the HapMap 3 European reference panel^41^. Furthermore, we used this test to explore if these SNPs are generally enriched for association with all (brain-related) phenotypes, or whether they exhibit some degree of outcome-specificity. For this purpose, we extended the LD-aware enrichment test to 21 additional traits for which GWAS results were available in the public domain. Some of the traits were chosen because they are phenotypically related to SZ (e.g., neuroticism, depressive symptoms, major depressive disorder, autism, and childhood IQ), while others were less obviously related to SZ (e.g., age at menarche, intracranial volume, cigarettes per day) or served as negative controls (height, birth weight, birth length, fasting (pro)insulin). The power of the LD-aware enrichment test primarily depends on the GWAS sample size of the target trait and results of our test would be expected to change as GWAS sample sizes keep growing.

**Supplementary Fig. 5** and **Supplementary Table 5.1** show the LD-aware enrichment of the 132 SNPs that are jointly associated with EA (*P_EA_* < 10^−5^) and SZ (*P_SZ_* < 0.05) across traits. First, we found significant joint LD-aware enrichment for SZ (P = 9.57 × 10^−66^), demonstrating that the genetic dependence between EA and SZ cannot be entirely explained by LD. We also found LD-aware enrichment of these SNPs for BIP, neuroticism, childhood IQ, and age at menarche. However, we found no LD-aware enrichment for other brain-traits that are phenotypically related to SZ, such as depressive symptoms, subjective well-being, autism, and attention deficit hyperactivity disorder. We also did not find LD-aware enrichment for most traits that are less obviously related to the brain and our negative controls. Furthermore, one of the novel SNPs we isolated shows significant LD-aware enrichment both for SZ and for BIP (rs7522116). The results suggest that the loci identified by the PPM are not simply related to all (brain) traits. Instead, they show some degree of phenotype specificity.

### Replication in the GRAS sample

We replicated the PPM analysis results in the GRAS sample (**Supplementary Note 10**) using a PGS that is based on the 132 independent EA lead SNPs that are also nominally associated with SZ (*P_EA_* < 10^−5^ and *P_SZ_* < 0.05, Supplementary Note 11). This PGS *(SZ132)* adds Δ*R*^2^ = 7.54% – 7.01% = 0.53% predictive accuracy for SZ case-control status to a PGS *(SZ_all)* derived from the GWAS on SZ alone (*P* = 1.7 × 10^−4^, **Supplementary Table 7.2.a**, Model 3).

### Prediction of schizophrenia measures in the GRAS patient sample

To explore the genetic architecture of specific SZ measures, we again used our replication sample (GRAS), which contains exceptionally detailed measures of SZ symptoms, severity, and disease history^4, 7, 28^. We focused on years of education, age at prodrome, age at disease onset, premorbid IQ (approximated by a multiple-choice vocabulary test), global assessment of functioning (GAF), the clinical global impression of severity (CGI-S), as well as positive and negative symptoms (PANSS positive and negative, respectively) among SZ patients (N ranges from 903 to 1,039, see **Supplementary Notes 10 and 12**). Consistent with the idea that EA is a predictor of SZ measures, our phenotypic correlations show that higher education is associated with later age at prodrome, later onset of disease, and less severe disease symptoms among SZ patients (**Supplementary Note 12, Supplementary Table 8.1** and **Supplementary Fig. 7**).

Our most direct test for genetic heterogeneity of SZ is based on PGS analyses that we performed using the detailed SZ measures among GRAS patients. If SZ is genetically heterogeneous, there is potentially relevant information in the sign concordance of individual SNPs with EA traits that may improve the prediction of symptoms (see **Supplementary Note 1** for formal derivations). We use a simple method to do this here: First, we construct a PGS for SZ that contains one SNP per LD-block that is most strongly associated with SZ. Overall, this score *(SZ_all)* contains 349,357 approximately LD-independent SNPs. Next, we split *SZ_all* into two scores, based on sign-concordance of the SNPs with SZ and EA. More specifically, one score contains all estimated SZ effects of SNPs that have concordant signs for both traits (174,734 SNPs with ++ or -- on both traits, *Concordant*) while the other contains the estimated SZ effects of the remaining SNPs with discordant effects (174,623 SNPs with +- or -+, *Discordant).* Note that splitting the *SZ_all* score this way is not expected to improve the prediction of symptoms if they share the same genetic architecture (i.e., if SZ was a genetically homogenous trait). We test this null hypothesis with an *F*-test that compares the predictive performance of models that include (i) the *SZ_all* and the EA score (*EA_all*) and (ii) the *Concordant, Discordant*, and *EA_all* scores (**Supplementary Note 1.3.2**). We also compare the performance of both of these models to a baseline that only includes the *SZ_all* score as a relevant predictor.

We found that the *EA_all* PGS is associated with years of education (*P* = 1.0 × 10^−6^) and premorbid IQ (*P* = 2.7 × 10^−4^) among SZ patients (**Supplementary Note 12** and **Table 2**). Consistent with earlier results^4^, we also found that none of the SZ measures can be predicted by the PGS for SZ (*SZ_all*, **Table 2**). However, splitting the PGS for SZ based on the sign-concordance of SNPs with EA *(Concordant* and *Discordant)* increased predictive accuracy significantly for severity of disease (GAF (*p_F_* = 0.023)) and symptoms (PANSS negative (*p_F_* = 0.007)) (**Table 2**). This increase in predictive accuracy is evidence for genetic heterogeneity of SZ (**Supplementary Note 1**). Specifically, our results indicate that SZ patients with a high genetic propensity for EA have better GAFs and less severe negative symptoms (PANSS negative). However, if the high genetic predisposition for EA is primarily due to loci that also increase the risk for SZ (i.e., high values on the *Concordant* score), this protective effect is attenuated. We repeated these analyses excluding patients who were diagnosed with schizoaffective disorder (SD, *N* = 198) and found similar results, implying that our findings are not only due to the presence of patients with SD (**Supplementary Note 12, Supplementary Table 8.4.a**).

We note that this implementation of our test for heterogeneity of SZ (**Supplementary Note 1**) is based on a conservative pruning algorithm that controls for LD both within and across the *Concordant* and *Discordant* scores. This limits the number of genetic markers in both of these scores, their expected predictive accuracy, and the power of the test. As an alternative, we also used a less conservative approach that only prunes for LD within scores, yielding 260,441 concordant and 261,062 discordant SNPs (**Supplementary Note 11.1.1**). Split scores based on this extended set of SNPs have higher predictive accuracy for all the SZ measures that we analysed (**Supplementary Table 8.8**), reaching Δ*R*^2^ = 1.12% (*p_F_* = 0.0004) for PANSS negative.

Finally, we show that randomly splitting the *SZ_all* score does not yield any gains in predictive accuracy (Supplementary Note 12 and Supplementary Table 8.5).

### Controlling for the genetic overlap between schizophrenia and bipolar disorder

The ongoing debate about what constitutes the difference between SZ and BIP^11–15^ suggests an additional possibility to test for genetic heterogeneity among SZ cases. While SZ and BIP share psychotic symptoms such as hallucinations and delusions, scholars have argued that SZ should be perceived as a neurodevelopmental disorder in which cognitive deficits precede the development of psychotic symptoms, while this is not the case for BIP^11–15^. However, cognitive deficits during adolescence are currently not a diagnostic criterion that formally differentiates SZ from BIP. As a result, many patients who are formally diagnosed with SZ did not suffer from cognitive impairments in their adolescent years, but their disease aetiology may be different from those who do. These differences in disease aetiology may be visible in how the non-shared part of the genetic architecture of SZ and BIP is related to measures of cognition, such as EA and childhood IQ.

We tested this by using genome-wide inferred statistics (GWIS)^42^ to obtain GWAS regression coefficients and standard errors for SZ that are “purged” of their genetic correlation with BIP and *vice versa* (yielding “unique” SZ_(min BIP)_ and “unique” BIP_(min sz)_ results, respectively). We then computed genetic correlations of these GWIS results with EA, childhood IQ, and (as a non-cognitive control trait) neuroticism using bivariate LD score regression^43^ and compared the results to those obtained using ordinary SZ and BIP GWAS results (**Supplementary Note 13**).

In line with earlier findings^8, 43^, we see a positive genetic correlation of ordinary SZ and BIP with EA. However, the genetic correlations between “unique” SZ_(min_bip)_ with EA and childhood IQ are negative and significant (*r_g_* = −0.16, *P* = 3.88×10^−04^ and *r_g_* = −0.31, *P* = 6.00×10^−03^, respectively), while the genetic correlation of “unique” BIP_(min SZ)_ with EA and IQ remain positive (*r_g_* ≈ 0.3) (**Fig. 3, Supplementary Table 9.2**). Thus, the slightly positive genetic correlation between SZ and EA^8, 43^ can be entirely attributed to the genetic overlap between SZ and BIP^42^, a result recently replicated using genomic structural equation modeling^44^. Overall, these results add to the impression that current clinical diagnoses of SZ aggregate over various non-identical disease aetiologies.

**Figure 3:**
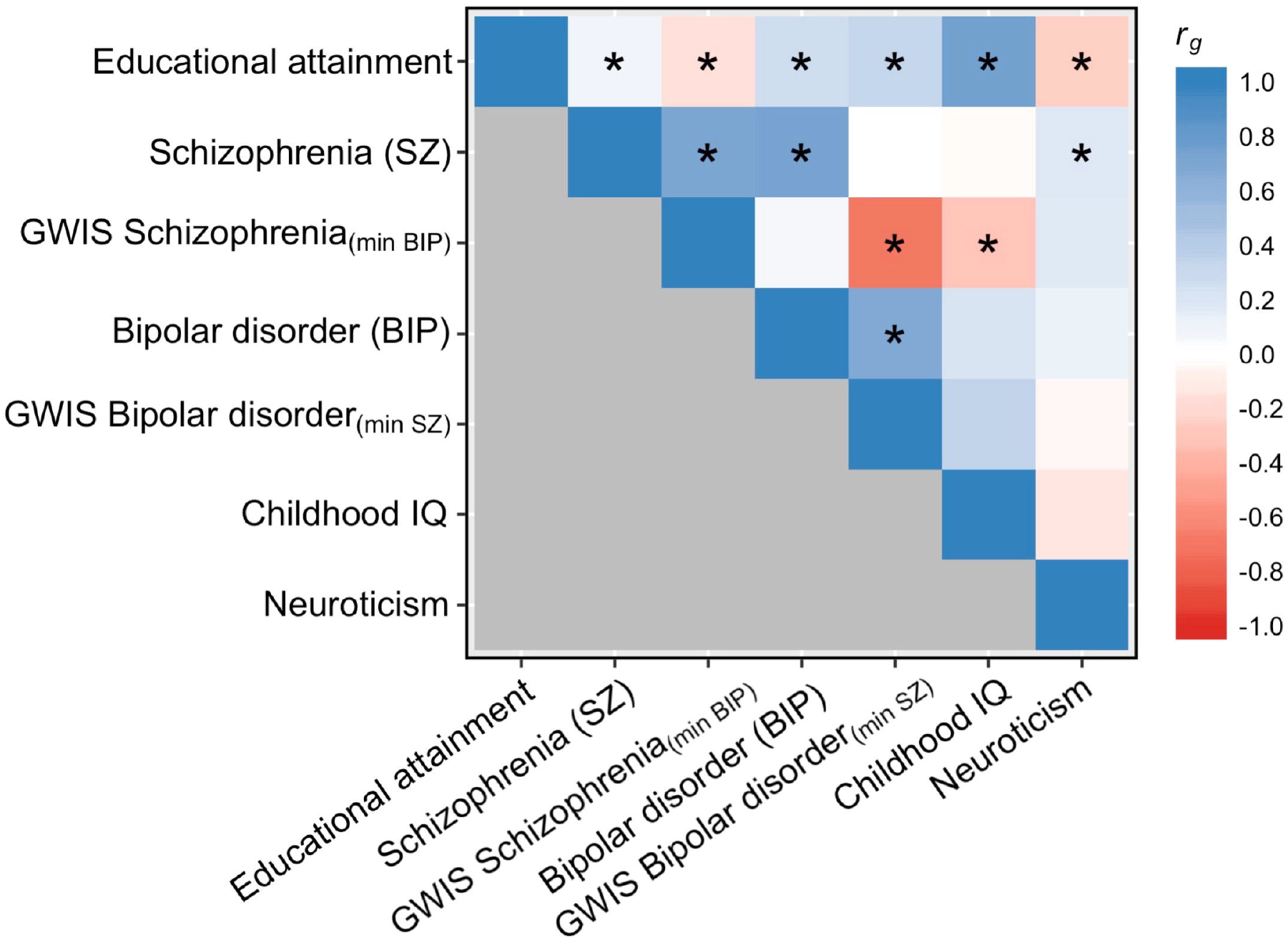
Genetic correlations of GWAS and GWIS results that are central to the relationship between SZ and EA. *Notes:* The heatmap displays the genetic correlations across 7 sets of GWAS or GWIS summary statistics. Genetic correlations were estimated with LD score regression.^43^ The colour scale represents the genetic correlations ranging from –1 (red) to 1 (blue). Asterisks denote significant genetic correlations at *P* value < 0.01.

Finally, simulations show that assortative mating is unlikely to be a major cause of the observed level of genetic dependence between EA and SZ (**Supplementary Note 14, Supplementary Fig. 9**).

## DISCUSSION

We explored the genetic relationship between EA and SZ using large, non-overlapping GWAS samples. Our results show that EA-associated SNPs are much more likely to be associated with SZ than expected by chance, i.e., both traits are genetically dependent. Overall, we isolated 21 genetic loci that are credibly associated with SZ by using EA as a proxy-phenotype, including two novel candidate genes, *FOXO6* and *SLITRK1.* Furthermore, we showed that EA GWAS results help to predict future GWAS findings for SZ in even larger samples.

Biological annotation of a broader set of SNPs that are jointly associated with EA (*P_EA_* < 10^−5^) and SZ (*P_SZ_* < 0.05) points to neurogenesis and synapse formation as potentially important pathways that may influence both traits.

However, the genetic loci that are associated with both traits do not follow a systematic sign pattern that would correspond to a strong positive or negative genetic correlation. Our follow-up analyses demonstrated that this pattern of strong genetic dependence but weak genetic correlation between EA and SZ cannot be fully explained by LD or assortative mating.

Instead, our results are most consistent with the idea that EA and SZ are both genetically heterogeneous traits that aggregate over various subphenotypes or symptoms with nonidentical genetic architectures. Specifically, our results suggest that current SZ diagnoses aggregate over at least two disease subtypes: One part resembles BIP and high IQ (possibly associated with *Concordant* SNPs), where better cognition may also be genetically linked to other BIP features such as higher energy and drive, while the other part is a cognitive disorder that is independent of BIP (possibly influenced by *Discordant* SNPs). This latter subtype bears similarity with Kraepelin’s description of dementia praecox^12^. Overall, our pattern of results resonates with the idea that cognitive deficits in early life may be an important differentiating factor between patients with BIP versus SZ psychosis.

Moreover, splitting the PGS for SZ into two scores based on the sign concordance of SNPs with EA enables the prediction of disease symptoms and severity from genetic data for the first time to some extent. We showed that this result is not driven by patients with SD and it cannot be repeated by randomly splitting the SZ score. Obviously, further replication of our results in other samples with high-quality SZ measures would be highly desirable.

The many sign-concordant loci that increase the risk for SZ but also improve the chance for higher education point to possible side-effects of pharmacological interventions that may aim to target biological pathways that are implicated by pleiotropic loci. Indeed, exploring pleiotropic patterns of disease-associated genes across a broad range of phenotypes (including social-scientific ones such as EA or subjective well-being^45^) may be a viable strategy to identify possible side-effects of new pharmacological products at early stages of drug development in the future.

Although the complexity of SZ remains astonishing, our study contributes to unravelling this complexity by starting at a genetic level of analysis using well-powered GWAS results. Our results provide some hope that a psychiatric nosology that is based on biological causes rather than pure phenotypical classifications may be feasible in the future. Studies that combine well-powered GWASs of several diseases and from phenotypes that represent variation in the normal range such as EA are likely to play an important part in this development. However, deep phenotyping of large patient samples will be necessary to link GWAS results from complex outcomes such as EA and SZ to specific biological disease subgroups.

## METHODS

A full description of all methods, materials, and results is available in the Supplementary Notes.

## GWAS

We obtained GWAS summary statistics on EA from the Social Science Genetic Association Consortium (<u>SSGAC</u>). The results are based on Okbay et al.^8^, including the UK Biobank. The PGC shared GWAS summary statistics on SZ with us that were reported in Ripke et al.^5^, but excluded data from our replication sample (GRAS, see **Supplementary Note 10**), yielding a total sample size of *n* = 34,409 cases and *n* = 45,670 controls. All cohorts that were part of both studies^5, 8^ were excluded from the meta-analysis on EA, yielding non-overlapping GWAS samples and *n_EA_* = 363,502. The original EA results file contained 12,299,530 genetic markers, compared to 17,221,718 in the SZ results file. We applied additional quality control steps: First, we excluded SNPs that were missing from large parts of both samples. Second, we excluded SNPs that were not available in both GWAS results files. Third, we excluded SNPs with non-standard alleles, mismatching effective alleles, and SNPs that exhibited strong differences in minor allele frequency in both results files. The remaining 8,240,280 autosomal SNPs were used in the proxy-phenotype and prediction analyses.

### Proxy-phenotype method (PPM)

We conducted proxy-phenotype analyses following a preregistered analysis plan (https://osf.io/dnhfk/), which specified that we would look up SZ results only for approximately independent SNPs with *P_EA_* < 10^−5^ in the independent EA sample. For LD-pruning in the EA GWAS results, we applied the clumping procedure in PLINK version 1. 9^46, 47^ using *r*^2^ > 0.1, a window of 1,000,000 kb, and the 1000 Genomes phase 1 version 3 European reference panel^48^.

### Biological annotations

To gain insights into possible biological pathways that are indicated by the PPM results, we applied DEPICT^8, 49^ using a false discovery rate threshold of ≤ 0.05. To identify independent biological groupings, we used the affinity propagation method based on the Pearson distance matrix for clustering^50^ (**Supplementary Note 8**).

### LD-aware enrichment of PPM results across different traits

For SNP *i* in trait *j*, we calculate the expected chi-square statistic as

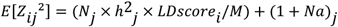

where *N* is the sample size of the target trait *j, h*^2^ is the heritability of trait *j*, 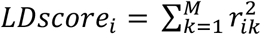 for SNP *i* is calculated using HapMap3 SNPs from European ancestry, *M* is the number of SNPs included in the calculation of the LD score (*n* = 1,173,569 SNPs), 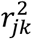 is the squared correlation between SNPs *j* and *k* in the HapMap3 reference panel, and 1 + *Na* is the LD score regression intercept for trait *j*. We calculated the LD score regression intercept and slope of the traits (*h*^2^) using LDSC^39^.

To determine whether a particular realization is significantly larger than expected (and thus the ratio 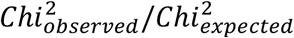 is significantly greater than one), we tested each particular observed *Z*–statistic (the square root of the *Chi*^2^) for SNP *j* against a normal distribution with variance 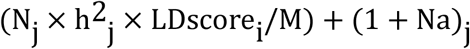.

### Replication of PPM results

We showed in our preregistered analysis plan that our replication sample (GRAS) is not large enough to replicate individual SNPs (https://osf.io/dnhfk/). Instead, we decided at the outset to attempt replication of the proxy-phenotype analysis results using a PGS that consists of the >80 most strongly associated, independent SNPs. The set that best meets this criterion are the 132 independent EA lead SNPs that are also nominally associated with SZ (<u>P_SZ_</u> < 0.05), see **Supplementary Note 6**. The PGS for this set of 132 candidate SNPs *(SZ_132)* was constructed in PLINK version 1.9^46, 47^ using the *β* coefficient estimates of the SZ GWAS meta-analysis.

### GWIS schizophrenia - bipolar disorder

To infer a SNP’s effect on SZ conditioned upon its effect on BIP, we approximated the following linear regression function:

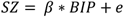

where the parameter *β* is estimated from the genetic covariance between SZ and BIP and the genetic variance in BIP as 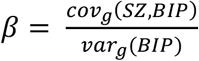. The residual (*e*) is actually our trait of interest, for which we use the term SZ_(min BIP)_. Using GWIS^42^, we inferred the genome-wide summary statistics for SZ_(min BIP)_ given the most recent PGC GWAS results for SZ (omitting the GRAS data collection)^5^ and BIP^51^. The effect size with respect to SZ_(min BIP)_ for a single SNP is computed as:

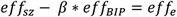

The standard error for each SNP effect is approximated using the delta method and accounts for the possible effect of sample overlap between the SZ and BIP GWAS.

As data input, we used the GWAS results on SZ (excluding the GRAS data collection) described in **Supplementary Note 3**. GWAS results for BIP^51^ (6990 cases, 4820 controls) were obtained from the website of the PGC (https://www.med.unc.edu/pgc/files/resultfiles/pgc.cross.bip.zip).

### Code availability

Source code for GWIS and LD-aware enrichment analyses will be made available through a GIT repository.

### Data availability

The GWAS summary statistics that were analysed during the current study are available on the website of the Social Science Genetic Association Consortium (SSGAC): http://www.thessgac.org/#!data/kuzq8. The GRAS data collection is not publicly available due to strict data protection laws in Germany for study participants that could potentially be identified. For further information, contact the study’s principal investigator Prof. Dr. Hannelore Ehrenreich.

## ACKNOWLEDGMENTS

This research was carried out under the auspices of the Social Science Genetic Association Consortium (SSGAC), including use of the UK Biobank Resource (application reference number 11425). We thank all research consortia that provide access to GWAS summary statistics in the public domain. Specifically, we acknowledge data access from the Psychiatric Genomics Consortium (PGC), the Genetic Investigation of ANthropometric Traits Consortium (GIANT), the International Inflammatory Bowel Disease Genetics Consortium (IIBDGC), the International Genomics of Alzheimer’s Project (IGAP), the CARDIoGRAMplusC4D Consortium, the Reproductive Genetics Consortium (ReproGen), the Tobacco and Genetics Consortium (TAG), the Meta-Analyses of Glucose and Insulin-related traits Consortium (MAGIC), the ENIGMA Consortium, and the Childhood Intelligence Consortium (CHIC). We would like to thank the customers and employees of 23andMe for making this work possible as well as Joyce J. Tung, Nick. A. Furlotte, and David. A Hinds from the 23andMe research team. This study was supported by funding from an ERC Consolidator Grant (647648 EdGe, Philipp D. Koellinger), the Max Planck Society, the Max Planck Förderstiftung, the DFG (CNMPB), EXTRABRAIN EU-FP7, the Niedersachsen-Research Network on Neuroinfectiology (N-RENNT), and EU-AIMS. Michel G. Nivard was supported by a Royal Netherlands Academy of Science Professor Award to Dorret I. Boomsma (PAH/6635). Additional acknowledgements are provided in the **Supplementary Note 16**.

## AUTHOR CONTRIBUTIONS

P.D.K. designed and oversaw the study and conducted proxy-phenotype analyses. V.B. and M.M. carried out analyses in the GRAS sample. R.V., C.A.P.B., M.N, and P.D.K. developed statistical methods. V.B. conducted bioinformatics and computed the LD-aware enrichment tests. C.A.P.B. and R.V. conducted simulation analyses. M.N. computed GWIS results, genetic correlations, and carried out pleiotropy analyses. R.K.L. assisted with biological annotation and visualization of results. P.D.K., V.B., M.M., and H.E. made especially major contributions to writing and editing. All authors contributed to and critically reviewed the manuscript.

## ADDITIONAL INFORMATION

**Supplementary Notes** are available at *Nature Communications’* website.

